# Adaptation strategies of iron-oxidizing bacteria *Gallionella* and Zetaproteobacteria crossing the marine–freshwater barrier

**DOI:** 10.1101/2024.02.28.582575

**Authors:** Petra Hribovšek, Emily Olesin Denny, Achim Mall, Håkon Dahle, Ida Helene Steen, Runar Stokke

## Abstract

Iron-oxidizing Betaproteobacteria and Zetaproteobacteria are generally associated with freshwater and marine environments, respectively. Despite repeated cross-environment observations of these taxa, there has been no focused exploration of genomes of marine *Gallionella* (Betaproteobacteria) to understand transitions between freshwater and marine habitats. Consequently, their roles in these environments remain uncertain. Here, we present strong evidence for co-occurrence of *Gallionella* and Zetaproteobacteria at deep-sea hydrothermal vents at the Arctic Mid-Ocean Ridges through metagenomic analyses. Phylogenomics analysis of *Gallionella* metagenome-assembled genomes (MAGs) suggests that seawater adaptation is an evolutionary event which occurred multiple times in distinct lineages. Similarly, several distinct evolutionary events for freshwater and terrestrial *Mariprofundus* and other Zetaproteobacteria are predicted. The presence of *cyc2* iron oxidation genes in co-occurring marine Betaproteobacteria and Zetaproteobacteria implies an overlap in niches of these iron-oxidizers. Functional enrichment analyses reveal genetic differences between marine MAGs of both iron-oxidizing groups and their terrestrial aquatic counterparts linked to salinity adaptation. Though scanning electron microscopy confirms the presence of Fe(III) oxyhydroxide stalks where *Gallionella* and *Mariprofundus* co-occur, *Gallionella* MAGs from hydrothermal vents lack evidence of putative stalk formation genes. *Mariprofundus* is therefore the likely sole stalk-producing iron-oxidizer in this environment. Conversely, discovery of putative stalk formation genes in *Mariprofundus* MAGs across the marine-freshwater barrier suggests that Fe(III) oxyhydroxide stalks might not be an exclusive signature for single iron-oxidizing taxa in marine and freshwater environments. Our research provides novel insights into the iron-oxidizing capacities, stalk production, environmental adaptation, and evolutionary transitions between marine and freshwater habitats for *Gallionella* and Zetaproteobacteria.

**Importance:** Iron-oxidizing bacteria play an important role in the global cycling of iron, carbon, and other metals. While it has previously been assumed that bacterial evolution does not frequently involve crossing the salinity barrier, recent studies indicate that such occurrences are more common than previously thought. Our study offers strong evidence that this also happens among iron-oxidizing bacteria, with new insights into how these bacteria adapt to the new environment, including hydrothermal vents and freshwater habitats. In addition, we emphasize the importance of accurate iron-oxidizing taxa identification through sequencing, rather than relying solely on the morphology of Fe(III) oxyhydroxides and environment. On a larger scale, microorganisms within established communities needing to respond to changes in salinity due to events like seawater intrusion in coastal aquifers underscore the importance of knowledge of transitions across habitat types with different salt concentration.

## 1 Introduction

Salinity is reported as the major factor shaping bacterial community composition between marine and freshwater environments (1). This is supported by numerous studies on communities along salinity gradients (2–5). Microbial marine–freshwater transitions are thought to be infrequent events, with marine and freshwater communities displaying differences in the abundance of major phyla and habitat-specific lineages (6). Conversely, many taxa are reproducibly observed in both environments, often at low abundances, supporting the idea that transitions between marine and freshwater ecosystems occur more frequently than anticipated (7). Several studies underscore the importance of knowledge of transitions across habitat types due to evidence of altered metabolic capacity of microbial communities across salinity gradients (8, 9). How taxa fulfilling critical ecosystem functions respond across marine and freshwater systems is key to document as salinity of water bodies continue to shift with changes in Earth climate regimes and anthropogenic disruption (10–12).

Organisms that move from freshwater to marine environments are exposed to osmotic stress caused by increased salt concentration in the environment. To adapt to osmotic pressure, marine microorganisms use strategies such as accumulation of ions using ion channels (13, 14) and import or production of compatible solutes intercellularly (15, 16). Transition to marine habitats involves acquiring genes linked to osmoregulation via horizontal gene transfer (9, 17, 18), often from marine bacteria (19, 20). In comparison to freshwater and terrestrial environments, microorganisms colonizing marine environments have been found to have smaller genomes (21, 22) and more acidic proteins (23–25).

Iron-oxidizing bacteria (FeOB) within the Betaproteobacteria family Gallionellaceae and the class Zetaproteobacteria are two groups for which genomic consequences of freshwater-marine transitions are unreported. FeOB play an important role in cycling of iron and other elements (26, 27) and heavy metal transport (Abramov et al., 2022). FeOB are early colonizers of metal surfaces, and promote corrosion (29, 30). Neutrophilic microaerophilic FeOB belonging to Betaproteobacteria within the family Gallionellaceae are usually observed in freshwater and other terrestrial environments (31–38). Before the discovery of Zetaproteobacteria, Fe(III) oxyhydroxide structures at deep-sea hydrothermal vents were mistaken to be produced by freshwater FeOB like *Gallionella* (39, 40). Iron-oxidizing Betaproteobacteria genera like *Gallionella, Leptothrix* and *Sideroxydans* (now placed under Betaproteobacteria within NCBI and Gammaproteobacteria within GTDB) are presently considered as limited to low-salinity environments and generally described as freshwater genera, while Zetaproteobacteria are described as marine (41). The present consensus is that Fe(III) oxyhydroxide stalks in freshwater can be attributed to Betaproteobacteria (*Gallionella*, *Ferriphaselus*) (42–46) and to Zetaproteobacteria (*Mariprofundus*) in marine environments (47, 48).

*Gallionella* has also been observed at some marine and brackish-associated sites (3, 49–53). Sequencing reads of iron-oxidizing genera of Gallionellaceae were also detected at deep-sea hydrothermal vents (54–61), with Zetaproteobacteria and Gallionellaceae FeOB often co-occurring. Similarly, Zetaproteobacteria are not only present at marine hydrothermal vents, but have also been found in coastal and brackish sediments (52, 62–70), microbial mats in estuarine environments (3), water columns of stratified estuaries (71, 72), and terrestrial aquatic environments (73–75), where they sometimes co-occur with Gallionellaceae (3, 28, 50, 76–83). Despite these observations of FeOB outside of their typical environments, their genomes have not been explored specifically to comprehend transitions between freshwater and marine habitats and their role in these environments remains unclear.

In this study, we present newly reconstructed metagenome-assembled genomes of *Gallionella* and *Mariprofundus* from hydrothermal vents. We show presence of *Gallionella* at 4% relative abundance at diffuse venting at Arctic Mid-Ocean Ridges, where scanning electron microscopy confirmed the presence of Fe(III) oxyhydroxide stalks where both *Gallionella* and *Mariprofundus* were detected. The co-occurrence of the two genera also implies an overlap in iron-oxidizing niches between Betaproteobacteria and Zetaproteobacteria. Our study gains insights into the iron-oxidizing capabilities, stalk production, adaptation to marine and freshwater environments, and the evolutionary transition between these habitats for FeOB genera *Gallionella* and *Mariprofundus*. Through genus-level pangenomes and functional enrichment analyses, we uncover differences in osmoprotectant biosynthesis and transporter genes, as well as specific trends in genome size, influencing the diversification of *Gallionella* and *Mariprofundus* across the marine-freshwater divide.

## 2 Results

### 2.0 *Gallionella* and Zetaproteobacteria co-exist at hydrothermal vents

Metagenome-assembled genomes (MAGs) representing two novel species or populations of *Gallionella* were reconstructed from iron oxide deposits, outer chimney wall and microbial iron mats (Fe mats) at diffuse hydrothermal venting sites at the Fåvne and Troll Wall vent fields (Fig 1, Fig 2 A-C, Table S1, Table S2). A single *Gallionella* MAG (AMOR20_M0989) was present with a coverage representing 4% of the total coverage of all detected MAGs in an iron oxide deposit at Fåvne (∼10 °C), while another *Gallionella* MAG (AMOR20_M0988) is present at 2% of the binned population in a chimney wall at Fåvne. Within these samples with highly abundant (1-4%) *Gallionella* MAGs, the iron oxide deposit also harbors several Zetaproteobacteria taxa (2 families and 6 defined genera as classified by GTDB) comprising 28% of the binned community altogether, while the chimney wall sample has co-occurring Zetaproteobacteria with 1% relative abundance (Fig 1a, Table S4). From the same metagenomes, 15 novel species-representative genomes of Zetaproteobacteria including several within genera *Mariprofundus* and *Ghiorsea* were recovered based on ANI analyses (95% cut-off) using publicly available MAGs (Table S3).

**Fig 1.**
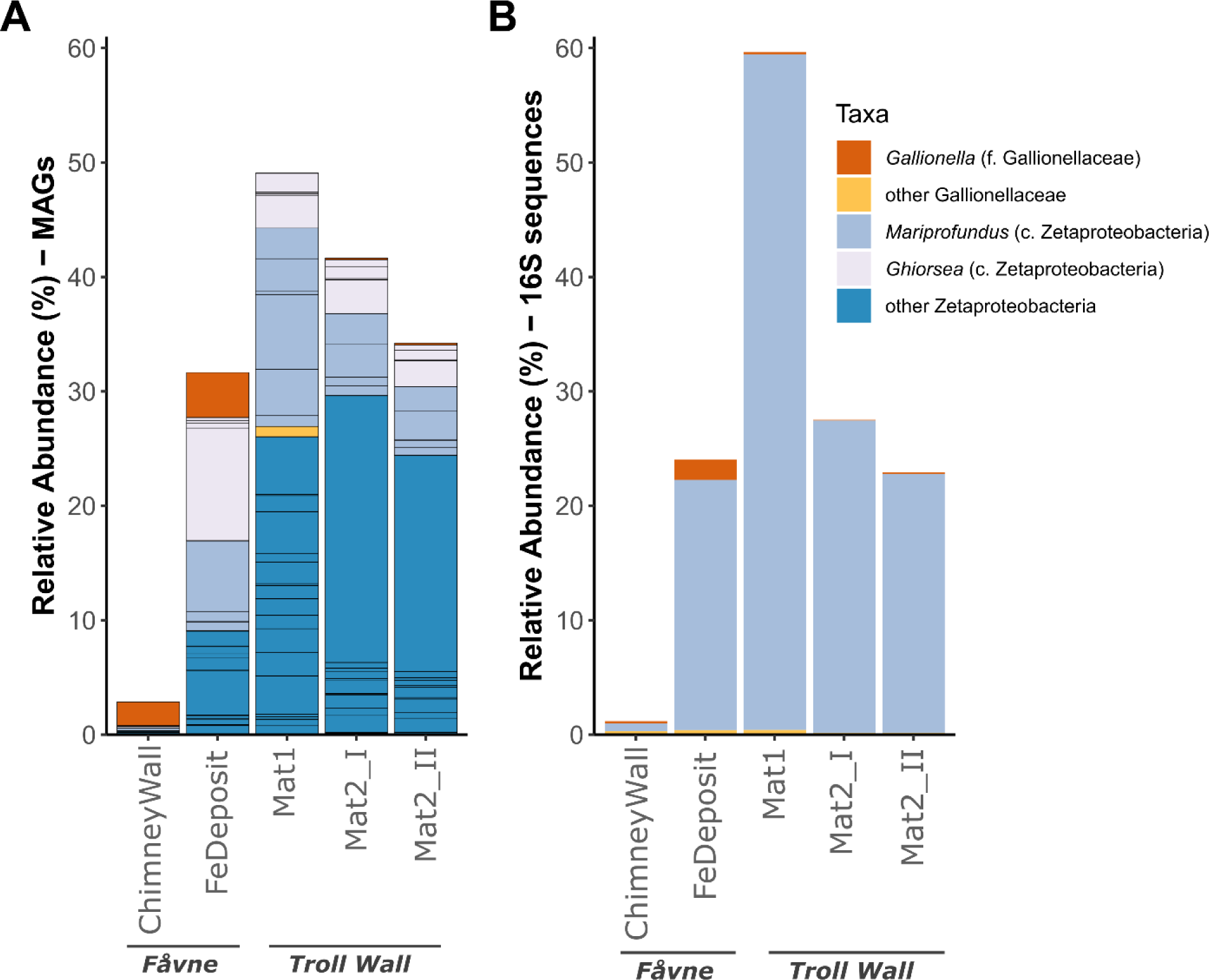
Diversity and relative abundance of FeOB at hydrothermal vents where Gallionella is detected at Fåvne and Troll Wall vent fields. A) Relative abundance of FeOB MAGs belonging to Gallionellaceae and Zetaproteobacteria. Relative abundances are based on reconstructed MAG coverage representing percentage of the total coverage of all detected MAGs in the sample. Black lines demonstrate the presence of several different MAGs within a taxonomic group, dereplicated at 98% ANI. B) Relative abundance of FeOB belonging to Gallionellaceae and Zetaproteobacteria based on all 16S sequences from the whole metagenome (PhyloFLASH).

**Fig 2.**
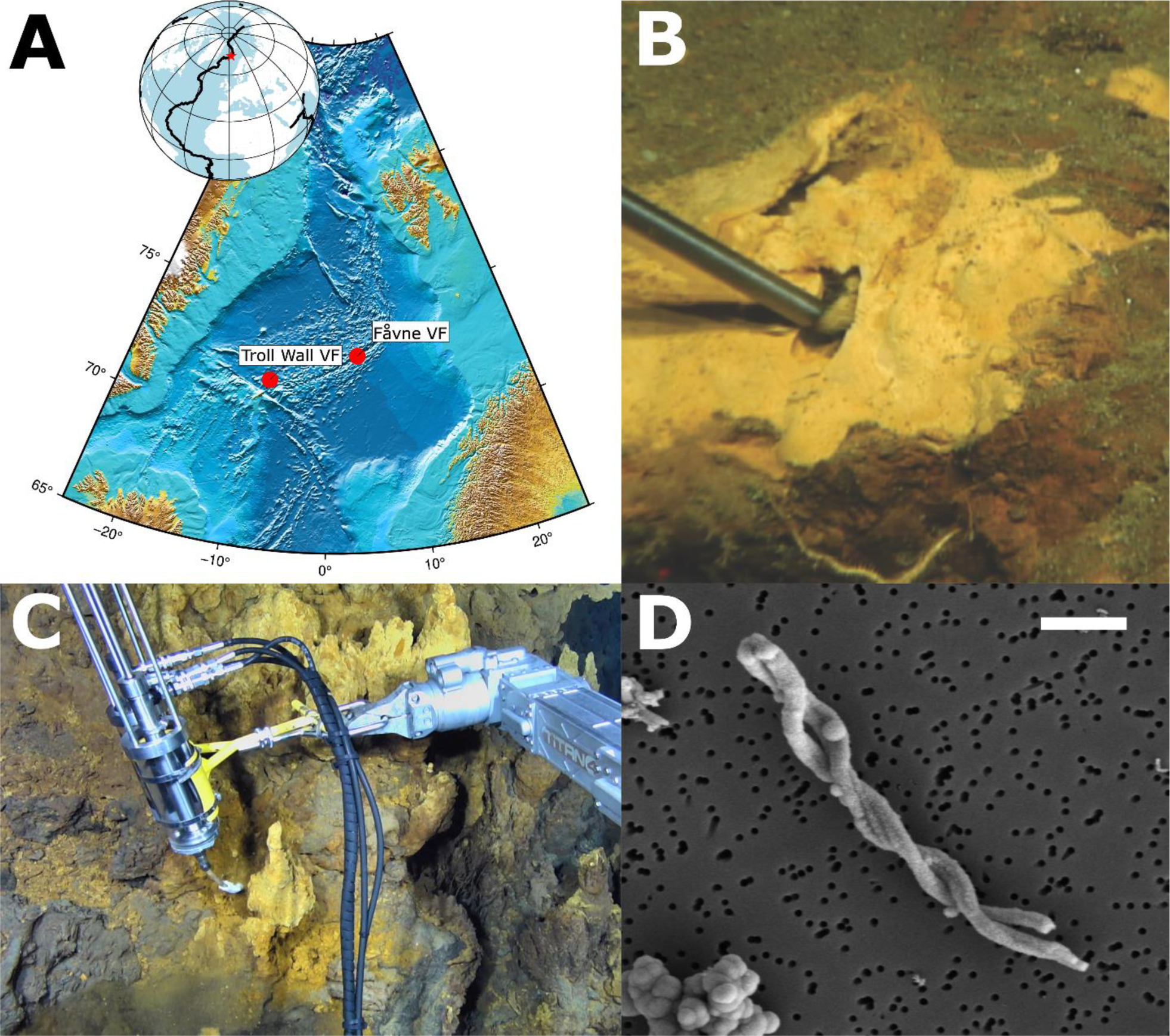
Hydrothermal vent sites where Mariprofundus and Gallionella co-exist. (A) Location of hydrothermal vent sites in the Arctic from where both Mariprofundus and Gallionella MAGs were reconstructed. Map generated using GMT and IBCAO grid. (B) Seafloor Fe mat at the Troll Wall vent field (Jan Mayen vent fields, Mat2 sample). Image modified from Vander Roost et al., 2017 under <u>Creative Commons Attribution License</u>. (C) Sampling of diffuse venting iron oxide deposit (∼10 °C) at Fåvne vent field. (D) Scanning electron microscopy showing biogenic Fe(III) oxyhydroxide stalks in iron oxide deposit at Fåvne, where both Mariprofundus and Gallionella MAGs were present. The scale represents 2 μm.

*Gallionella* MAGs are present at lower abundances at Troll Wall in comparison to Fåvne. In the Troll Wall vent Fe mats (between 2.5 and 5 °C) where a Gallionellaceae MAG is present at 1% (Table S5), Zetaproteobacteria MAGs are present at 48% of the binned community. When considering MAGs with coverage values above 0.5, *Gallionella* MAGs co-occur with Zetaproteobacteria in all samples taken from the Fåvne and Troll Wall vent fields (Table S4). The diversity of FeOB Gallionellaceae is lower than for Zetaproteobacteria, also at the genus level (Fig 1).

Scanning electron microscopy confirms the presence of Fe(III) oxyhydroxide stalks in the iron oxide deposit sample where both *Gallionella* and *Mariprofundus* MAGs were detected (Fig 2 D). In *Gallionella* MAGs from Fåvne and Troll Wall vent fields, genes potentially needed for stalk formation (*sfz1-4*) were not identified (Fig 3). *Gallionella* MAGs sourced from other marine sites follow a similar trend. As a follow-up to this finding, an assessment of all Zetaproteobacteria MAGs stalk formation potential revealed *Mariprofundus* MAGs from freshwater environments do possess putative stalk formation genes (Fig 4).

**Fig 3.**
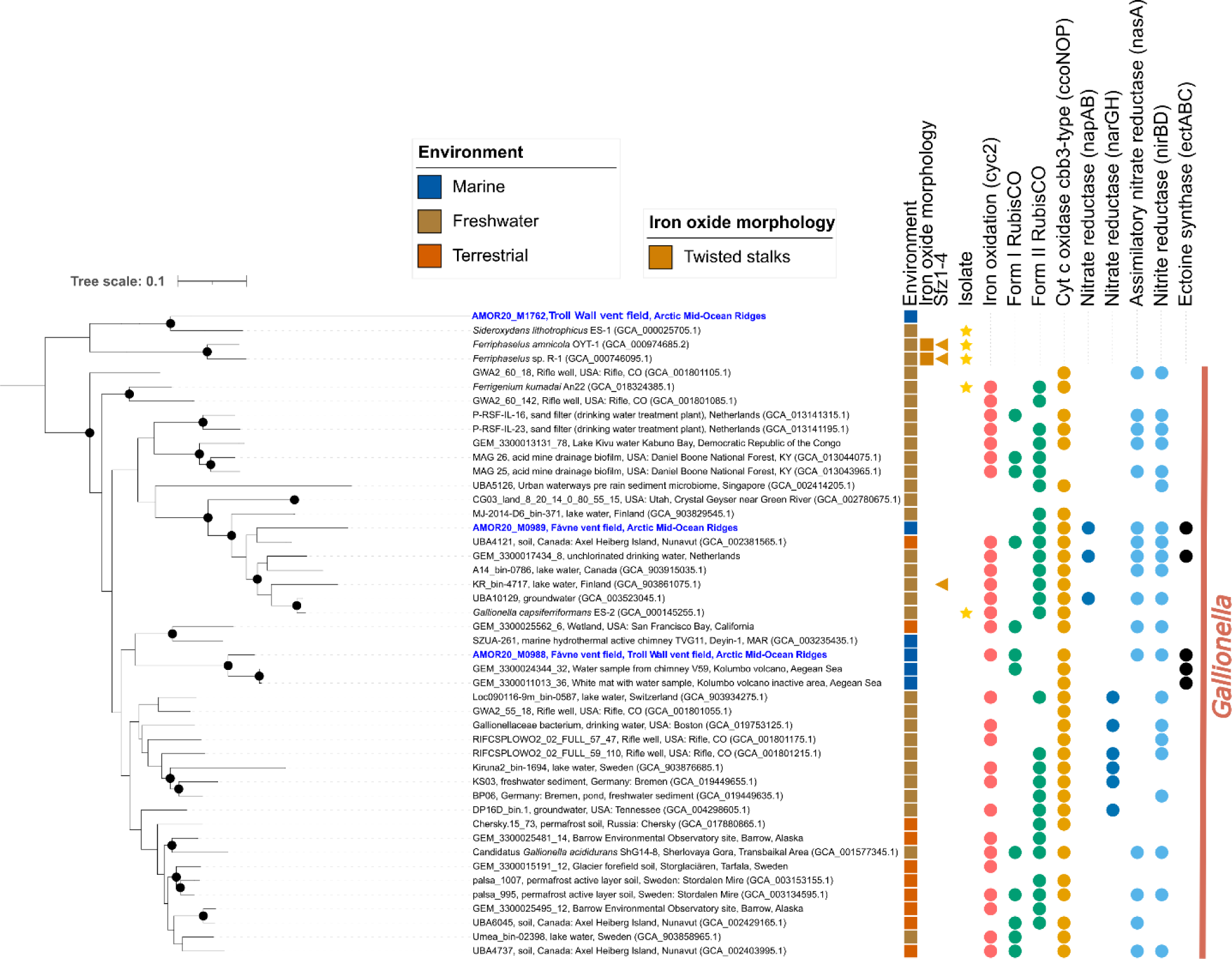
Phylogeny of Betaproteobacteria within the genus Gallionella. The phylogenomics tree is based on a concatenated alignment of a manually curated set of 15 single copy gene markers (Table S6) using MAGs from this study and references. All MAGs represent a species-level cluster (using 95% ANI cutoff), except for marine Gallionella MAGs, which were not clustered. Sfz1-4: potential stalk formation genes. Environmental assignments are based on NCBI metadata. Completeness and contamination values for each MAG are based on CheckM2 predictions. Genomes reconstructed from the Fåvne vent field and Troll Wall vent field are shown in blue. The maximum likelihood tree was constructed with IQTREE using substitution model GTR20+F+R6. Black node circles mark branches with support values higher than 80% with SH-like approximate likelihood ratio test and 95% with ultrafast bootstrapping, both including 1000 iterations. The tree is rooted using non-Gallionella Gallionellaceae MAG sequences as an outgroup.

**Fig 4.**
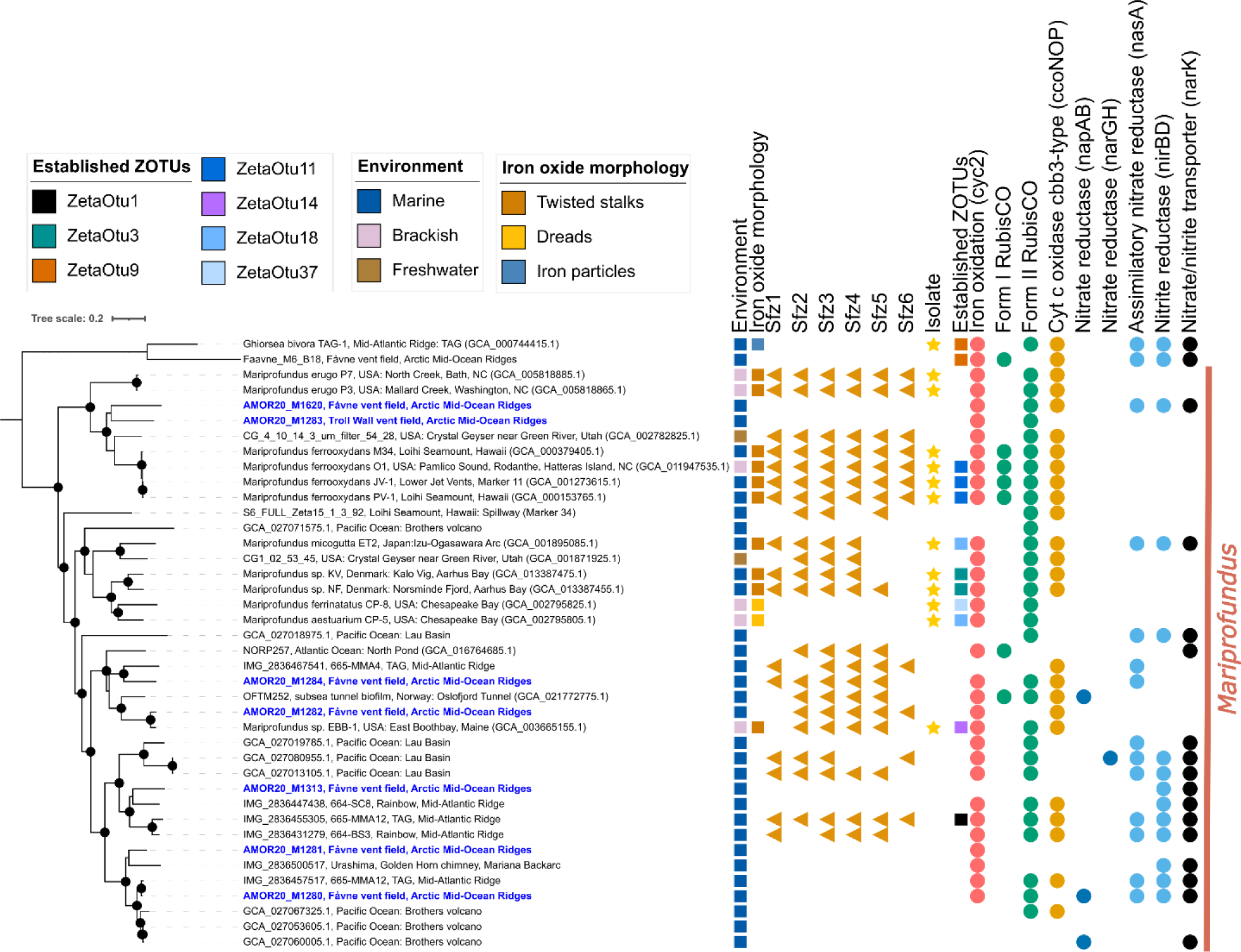
Phylogeny of Zetaproteobacteria within the genus Mariprofundus. The phylogenomics tree is based on a concatenated alignment of a manually curated set of 12 single copy gene markers (84) using MAGs from this study and references. Sfz1-6: potential stalk formation genes in Zetaproteobacteria. The environments specified are based on NCBI metadata. Genomes shown in blue have been reconstructed from the Fåvne vent field and Troll Wall vent field. The maximum likelihood tree was generated using IQTREE with substitution model LG+F+R9. Black node circles mark branches with support values higher than 80% with SH-like approximate likelihood ratio test and 95% with ultrafast bootstrapping, both including 1000 iterations. The root of the tree is based on Ghiorsea MAG sequences as an outgroup.

### 2.1 Marine *Gallionella* are closely related to their freshwater counterparts

To assess whether marine *Gallionella* populations are closely related to freshwater *Gallionella*, we performed phylogenomic analysis (Fig 3). The two recovered deep-sea hydrothermal vent *Gallionella* MAGs from Fåvne and Troll Wall are AMOR20_M0989 (97.1% completeness, 0.1% contamination), with closest relative from permafrost (GEM_3300012005_15, 74.8% ANI), and AMOR20_M0988 (93.8% completeness, 4.9% contamination), with closest relative from a hydrothermal system of Kolumbo volcano (GEM_3300011013_36, 84.4% ANI). *Gallionella* genome SZUA-261 from active and inactive marine hydrothermal chimneys at the Deyin-1 vent on the Mid-Atlantic Ridge (57) is different from both AMOR *Gallionella* MAGs at 70% ANI. Phylogenomic analysis using 15 manually curated marker genes suggests two separate evolutionary events for marine *Gallionella*, which cluster together with their freshwater and terrestrial counterparts (Fig 3, Fig S1). The presence/absence dendrogram in our pangenome also shows 2 different marine *Gallionella* clusters. In contrast, phylogenomic analysis using different sets of markers shows either three or two clusters of the marine *Gallionella* (Fig S2, Fig S3). Four marine *Gallionella* MAGs are unique species-level representative genomes, with two MAGs from subsea Kolumbo volcano sharing more than 98% ANI similarity.

### 2.2 Marine Zetaproteobacteria are closely related to their freshwater counterparts

To assess to what extent close evolutionary relationships can be found between Zetaproteobacteria in marine and freshwater environments, phylogenomics including all AMOR and publicly available Zetaproteobacteria MAGs was performed. Genome-resolved metagenomics of samples from the Troll Wall vent field resulted in 16 Zetaproteobacteria MAGs in addition to the already published MAGs from Fe mats on Fåvne vent field black smokers (84) (Table S3). Phylogenomic analysis revealed two distinct clusters of freshwater *Mariprofundus* grouping among marine representatives (Fig 4) Similarly, several distinct clusters of hot spring Zetaproteobacteria were clustering together with their marine counterparts (Fig S4), suggesting adaptations of marine Zetaproteobacteria to freshwater conditions is an evolutionary event that has occurred multiple times in distinct lineages.

### 2.3 Iron and carbon metabolism is conserved in *Gallionella* in the marine environment

Analysis of iron oxidation genes shows a conserved Fe energy metabolism in marine *Gallionella* at the Fåvne and Troll Wall vent fields. Several *Gallionella* MAGs belonging to a cluster of marine *Gallionella* encode a *cyc2* gene, while in individual marine *Gallionella* MAGs *cyc2* genes were not detected but are present in closely related freshwater and terrestrial MAGs (Fig 3). Incomplete aerobic and anaerobic respiratory pathways are present in marine *Gallionella*, as well as genes involved in CO_2_ fixation. A cluster of several marine *Gallionella* MAGs encode a Form I Rubisco adapted to higher O_2_ concentrations, while the individual marine *Gallionella* MAG encodes a Form II Rubisco adapted to low oxygen concentrations (85, 86). Marine *Gallionella* MAGs also contain genes for cbb3-type cytochrome oxidase; an oxidase better adapted to low oxygen conditions (87). To further assess the ability of *Gallionella* and Zetaproteobacteria to oxidize iron outside their typical environments, MAGs from various global locations were included. Iron oxidation genes are present in FeOB genomes across the marine-freshwater barrier, with *cyc2* also present in freshwater Zetaproteobacteria MAGs (Fig 4, Fig S4). No uptake hydrogenases indicating hydrogen as a putative alternative electron donor were detected in *Gallionella* MAGs from Fåvne vent field or other marine environments.

### 2.4 Environmental adaptation in marine *Gallionella* genomes

Functionally enriched genes in marine *Gallionella* are associated with environmental adaptation, such as salinity adaptation using antiporters and osmolyte production, heavy metal resistance, resistance against viral attack and other organisms, uptake of potential nutrients or antiporters, transport systems and membrane-bound proteins (Table S7). Sulfatase genes are enriched and present in 3 MAGs of marine *Gallionella.* Enriched genes in marine *Gallionella* MAGs encode for biosynthesis of osmolyte ectoine (88) and biosynthesis of 9-membered enediyne core, known for antibiotic and antitumoral activities (89). Genes involved in trehalose biosynthesis are enriched in terrestrial MAGs. Genes for potassium transporter are enriched in freshwater and terrestrial *Gallionella* MAGs (more than 80% of MAGs), while genes for sodium/alanine symporter, sodium/proline symporter, sodium/dicarboxylate symporter and sodium/hydrogen antiporter are enriched in marine *Gallionella* MAGs.

The ectoine synthase gene was present in a majority of marine *Gallionella* (4 out of 5 MAGs) (Fig 3, Fig 5). Exploring the potential acquisition of ectoine synthase genes through horizontal gene transfer, BLAST searches and phylogenetic tree reconstruction pinpoints potential sources such as other Gallionellaceae members, unknown Gammaproteobacteria, and Gammaproteobacteria of genus *Rugosibacter* and *Microbulbifer,* commonly present in marine sediments (90) (Fig 5, Fig S5). Closest ectoine synthase gene relatives to marine *Gallionella* within Gallionellaceae are *Gallionella* and *Sideroxydans* from brackish environments and a non-iron oxidizing *Nitrotoga* sp. BS from a wastewater treatment sample tolerant to up to 1% salt (91). Within the *Gallionella* genus, ectoine synthase genes exhibit 80-85% identity, and 60-65% similarity to genes of *Microbulbifer* found in marine and brackish environments. Among these genes, *ectB* and *ectC* demonstrate a more conserved trend compared to *ectA* (Fig 5).

**Fig 5.**
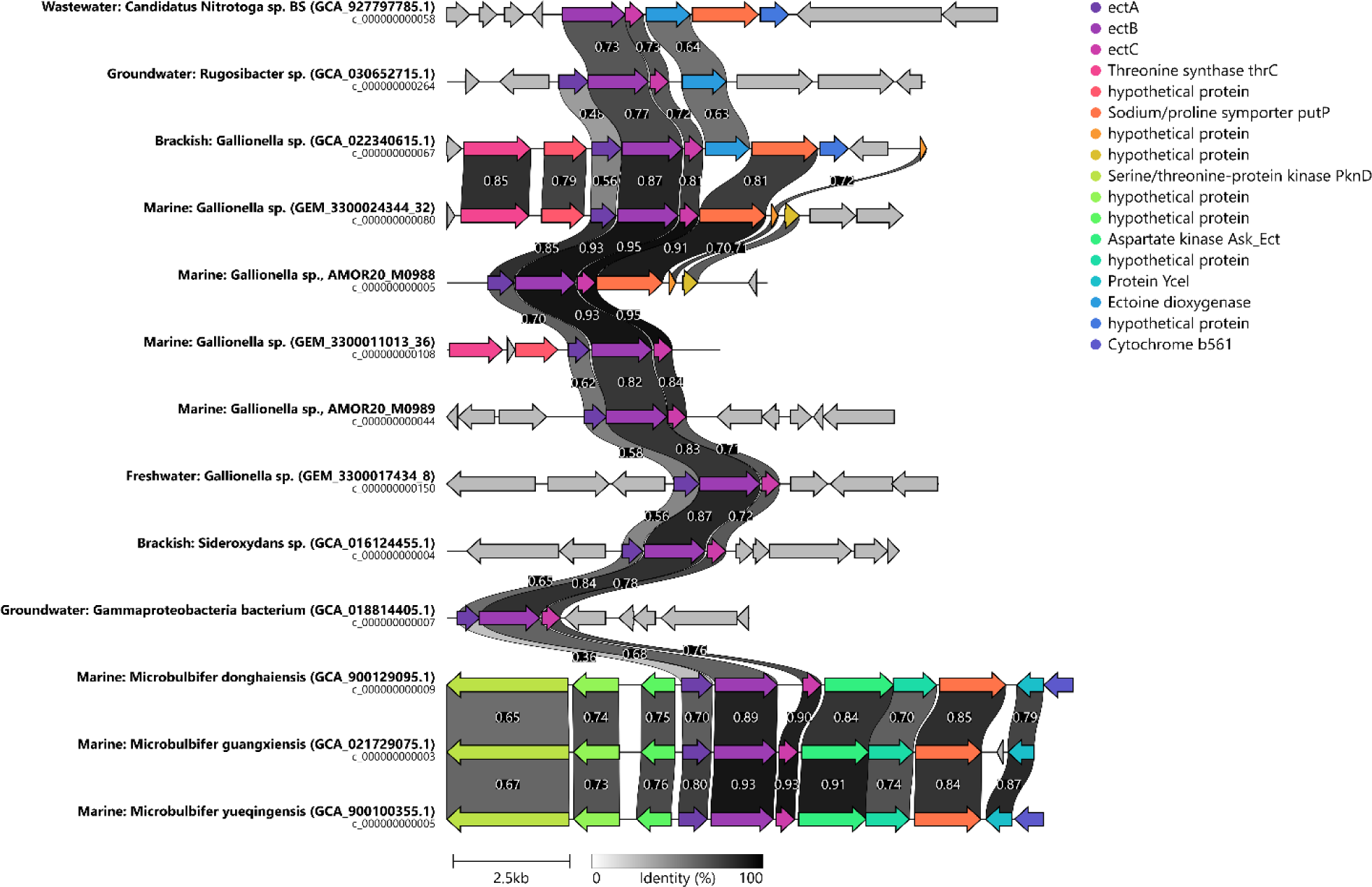
Gene neighborhood of ectoine synthase genes present in Gallionella MAGs including gene sequences from all publicly available Gallionella genomes and closely related and publicly available reference genomes. Ectoine synthase genes (ectA, ectB, ectC) and flanking genes are highlighted by colors as in the legend.

### 2.5 Environmental adaptation and nitrate assimilation in Zetaproteobacteria genomes

Genes for potassium/hydrogen antiporter that regulates potassium levels within cells, preventing toxicity in low osmolarity situations (92, 93) are enriched in more than 66% of freshwater *Mariprofundus* MAGs and ca 10% marine MAGs, while genes for sodium/hydrogen antiporter are enriched in brackish and marine Zetaproteobacteria MAGs. Moreover, genes for sodium or hydrogen/acetate symporter are enriched both in marine and freshwater Zetaproteobacteria MAGs. Twenty-five percent of Zetaproteobacteria MAGs assigned as brackish (8 MAGs) and a few marine MAGs possess genes responsible for producing osmoprotectants like trehalose, a spermidine/putrescine transport system, acetoin utilization protein and glutamate/leucine dehydrogenase. *NarK* gene encoding for a nitrate/nitrite transporter (94, 95) possibly involved in nitrate assimilation, is exclusively present in 58% of marine MAGs (Table S8). A similar trend is observed in the genus *Mariprofundus* (Table S9). Mercury resistance gene (*merR1*) is found in 15 Zetaproteobacteria MAGs exclusively from hydrothermal vents (Fig S4).

### 2.6 Further outlook into genomic adaptations

Generally small average genome size (2.19±0.16 Mb), high percentage of coding density (91±2.1 %) and low GC content (49.7±2.5 %) are observed in marine *Gallionella* MAGs (Table S10). Performing both the one-way ANOVA and the Kruskal-Wallis test with appropriate post hoc tests to examine the effects of the environment on genome size, differences were detected between marine and terrestrial *Gallionella* genomes. Marine genomes of Zetaproteobacteria are smaller compared to their brackish and freshwater counterparts, while marine *Mariprofundus* genomes are smaller than freshwater ones (Fig 6, Table S10). Genome size varies greatly within environment groups, however, and due to few genomes in some groups such as the marine *Gallionella* genomes, the power analysis indicates that sampling of more genomes is required to be confident in the conclusion at significance level of 0.05.

**Fig 6.**
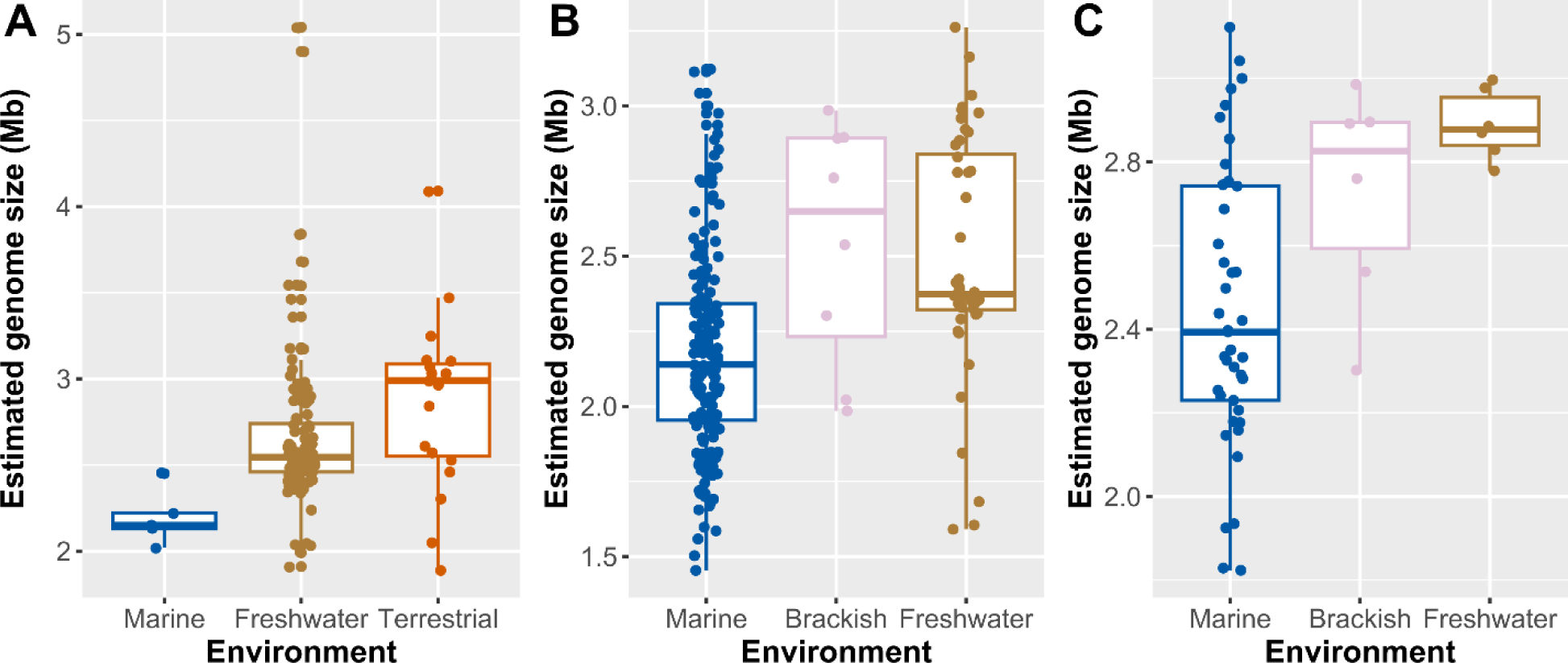
Estimated genome sizes of Gallionella and Zetaproteobacteria genomes from marine, brackish, freshwater, and terrestrial environments. Publicly available genomes and MAGs from this study. Boxplots visualize the distribution of estimated genome sizes for each environment, showing mean and standard deviation with points visualizing estimated genome sizes of all individual MAGs. A) Gallionella MAGs. B) Zetaproteobacteria MAGs. C) Mariprofundus MAGs.

Whole-proteome isoelectric point (pI) comparison of *Gallionella* predicts proteomes exhibit mostly similarities across the marine-freshwater barrier (Fig S6), while differences between environments are more obvious in Zetaproteobacteria and *Mariprofundus* (Fig S7, Fig S8). Average relative frequencies for acid, neutral and basic pIs show little difference between environments (Table S11). As with genome sizes, sampling of more genomes is required for statistical analyses with great confidence.

## 3 Discussion

This comparative genomics study investigated how members of freshwater *Gallionella* may adapt to marine conditions and how marine Zetaproteobacteria may adapt to freshwater conditions. Our study includes the reconstruction of novel *Gallionella* MAGs from diffuse venting at marine hydrothermal vents, providing further evidence that the adaptive transition from a freshwater setting to a marine setting, or vice versa, can occur over relatively short evolutionary timescales among *Gallionella*. Here, we consider aspects of the transition of the marine-freshwater barrier and offer a model involving adaptations of FeOB to higher salinity via horizontal gene transfer and possible genome reduction.

### 3.0 Autotrophic and iron oxidizing *Gallionella* and Zetaproteobacteria co-exist in hydrothermal vents

Earlier investigations of low salinity environments revealed the co-occurrence of Zetaproteobacteria and Gallionellaceae FeOB, in settings including brackish waters, terrestrial hot springs, benthic microbial mats, and fjord sediments (3, 28, 50, 76–83). In marine hydrothermal systems, co-existence of Zetaproteobacteria and Gallionellaceae FeOB was also indicated in iron mats, iron mounds, and active and inactive hydrothermal chimneys (54–58, 60). In addition to *Gallionella*, closely related iron-oxidizing Betaproteobacteria like *Sideroxydans* and *Leptothrix* have been detected with 16S rRNA sequences at the Troll Wall hydrothermal vent field (56).

*Gallionella* and Zetaproteobacteria MAGs recovered from the same diffuse hydrothermal vent samples at AMOR provide further evidence that co-existence of the two FeOB groups is globally widespread, with Zetaproteobacteria consistently present wherever *Gallionella* is detected in the marine environment. The only exception are the inactive sulfide chimneys from East Pacific Rise where only *Gallionella* sequences were detected (59). *Gallionella* and Zetaproteobacteria are predicted to oxidize iron regardless of environment, implying an overlap in iron-oxidizing niches between Betaproteobacteria and Zetaproteobacteria within the same community. Marine *Gallionella* present at such high abundances (up to 4% of the community) at Fåvne vent field may indicate that *Gallionella* FeOB are responsible for significant iron oxidation at diffuse venting zones of this hydrothermal field. *Gallionella* and Zetaproteobacteria at hydrothermal vents are possibly both involved in chemolithoautotrophy and processes contributing to the cycling of iron and carbon in the same environment. Interactions between these seemingly functionally redundant FeOB, such as competition, remain to be studied.

### 3.1 Identity of FeOB forming Fe(III) oxyhydroxide stalks in marine and freshwater environments

Stalks of *Gallionella* and *Mariprofundus* are remarkably similar (96). Stalk-producing representatives of both Betaproteobacteria and Zetaproteobacteria were previously found to possess the *sfz* gene cluster, putatively involved in stalk formation (44, 97). Stalks and the *sfz1-4* genes (97) are suggested to be limited to the genera *Gallionella* and *Ferriphaselus* within Gallionellaceae (46). Based on the presence of both *Mariprofundus* and *Gallionella* in hydrothermal iron oxide deposits, observed stalks could theoretically be produced by either *Mariprofundus* or *Gallionella*. Even so, no evidence of putative stalk formation genes is found in *Gallionella* MAGs at hydrothermal vents (Fig 3), whereas co-occurring *Mariprofundus* MAGs contain *sfz* genes (Fig 4). It is therefore more likely that stalks are produced by *Mariprofundus* in this environment. *Sfz* genes do not seem widespread in *Gallionella* populations, regardless of habitat, as previously observed (46), but are rather more widely distributed in *Mariprofundus* populations. *Mariprofundus* genomes sourced from diverse environments spanning marine, brackish, and terrestrial hot springs, on the other hand, have putative stalk formation genes (Fig 4). Consequently, Fe(III) oxyhydroxide stalks are not necessarily a definitive signature for *Mariprofundus* and *Gallionella* in marine and freshwater environments, respectively. Identity of FeOB groups has often been interpreted solely on the morphology of Fe(III) oxyhydroxides and the environment (40, 98–103). We emphasize that DNA-based identification methods are necessary for accurate FeOB identification rather than reliance on stalk morphology and environment alone. It remains unclear whether marine *Gallionella* and freshwater *Mariprofundus* express these genes and produce stalks; only microscopy and cultivation efforts will show whether this is the case. Not all *Gallionella* isolates produce stalks (45, 104, 105), neither do all *Mariprofundus* isolates (70, 97, 106). Even when a FeOB species has the ability to produce stalks, they don’t appear to be essential for growth (107), as stalk formation might be affected by conditions such as cell number, Fe(II) concentrations and pH (97, 104).

### 3.2 FeOB seem to have crossed the marine-freshwater barrier in several evolutionary events

Capacity to thrive in diverse salinity conditions is proposed to have developed early in bacterial evolution (8). Although marine and freshwater bacteria co-occur in brackish environments, true microbial freshwater-marine transitions are assumed as infrequent phenomena due to the necessity to adjust to drastic changes in physicochemical conditions. Habitat transitions may result in differences in proteomes and potentially substantial changes in central metabolism, which suggest significant evolutionary time has passed for the salinity adaptation to occur (23). General observation also supports transitional infrequency as dominant freshwater and marine taxa are usually not closely related (6). Upon close examination, it appears that certain transitioning microorganisms may in fact be present within the “rare biosphere” (7), including *Gallionella* and Zetaproteobacteria.

Our phylogenetic analyses suggest that FeOB transitions between freshwater and marine environments could have occurred several times, with several *Gallionella* species colonizing the marine environment independently. Marine *Gallionella* genomes appear closely related to freshwater *Gallionella* (Fig 3), confirming previous findings based on 16S rRNA gene sequences (56). As *Gallionella* seems widely present in marine environments, they appear to have successfully transitioned across the freshwater–marine boundary. Similarly, multiple crossings of the marine-freshwater barrier are observed within Zetaproteobacteria, including the genus *Mariprofundus* (Fig 4, Fig S4). These results imply complex evolutionary histories which could be improved in resolution with more genomes and updated 16S rRNA gene sequence analyses. The direction of adaptation (fresh to saline, or vice versa) remains unclear for both groups of FeOB. Reconstructing habitat of origin for FeOB lineages might be aided through phylogenomic analyses in tandem with phylogenetic analysis of genes involved in synthesis of osmolytes, which has proved instrumental for similar investigations in other taxa (108).

Transition of FeOB between freshwater and marine environments could have been promoted by adaptations in brackish environments as hypothesized for some other microorganisms crossing the barrier (109), though no substantial proof of such an event was detected from our phylogenomic reconstructions. In brackish microorganisms, both freshwater and marine characteristics (salt adaptation mechanisms) are present (93). *Gallionella* have ectoine synthase genes which are closely related to one another. Possible sources of the ectoine synthase genes may be through organism(s) in brackish or freshwater settings already adapted to conditions within a marine/estuarine environment or through interactions in groundwater exposed to inflow of saltwater. The combined evidence of phylogenetically similar *Gallionella* genomes across geographically spread marine and freshwater settings, likely horizontal gene transfer, and lack of pronounced differences in proteome isoelectric profiles, suggests that *Gallionella* freshwater–marine transition might have been driven by a rapid rather than gradual adaptation, as suggested for several other taxonomic groups (110).

### 3.3 Possible adaptations of *Gallionella* to marine environment

Comparing *Sideroxydans* and *Gallionella* based on genomic and ecological analysis, *Sideroxydans* has been seen as perhaps better adapted to conditions with higher salinity than *Gallionella* (3). In this study with observations of highly abundant marine *Gallionella* species, we now know *Gallionella* can also be adapted to higher salinity. Salinity of water masses above Mohns Ridge is expected to be 34.9 ppt – very close to the global average of 35 ppt, based on vertical CTD profile data (111). Given the high degree of dilution of hydrothermal fluids with ambient seawater upon exit at the seafloor (typically >80 to 90%), and the relatively limited range of salinity in most endmember vent fluids (112), most low temperature/diffuse vent fluid salinities do not differ substantially from background saltwater salinity (113). For this reason, we argue that *Gallionella* at hydrothermal vents on the AMOR are likely only exposed to salinities comparable to seawater, and we consider the possibility of some cryptic microniches of low salinity at these settings highly unlikely. This implies therefore that some *Gallionella* species are fully adapted to marine environments.

There are certain bioenergetic costs associated with all strategies of adaptation to higher salinity, which come with evolutionary speculations. Accumulation of salts inside the cell though requires the intracellular cell machinery to be adapted to higher salt concentrations (114), while this is not required by the compatible solute strategy which may not require a such a large modification to the proteome (15). However, such microorganisms need to instead invest energy in ion pumps to maintain low ionic concentrations in the cell.

Salinity seems to play a role in the adaptation of marine *Gallionella* (Fig 7). Genes for synthesis of ectoine, a compatible solute, are enriched in marine *Gallionella* genomes (Fig 2). Genes for synthesis of compatible solutes like ectoine, possibly obtained via horizontal gene transfer, allow *Gallionella* in the marine environment to overcome osmotic stress and adapt to environments of different salinities. Production of the compatible solute ectoine as an adaptation is seen in several marine and halophilic microorganisms (88, 115–117). All marine *Gallionella* and other Gallionellaceae that have ectoine synthase genes they potentially acquired from Gammaproteobacteria. *Gallionella* genomes from different environments show differential presence of several ion pump genes that typically participate in establishing ion gradients across the cellular membrane. Freshwater organisms have been found to use ion/potassium channels, as opposed to ion/sodium channels used by marine organisms (22, 93), which was also evident in *Gallionella* MAGs. The differential adaptation to the environment with elevated salinity also involved different genes involved in lipid and fatty acid metabolism (93, 118). Although not exclusively observed in marine *Gallionella* MAGs, the production of extracellular polymeric substances might also assist in adapting to elevated salinity levels (119).

**Fig 7.**
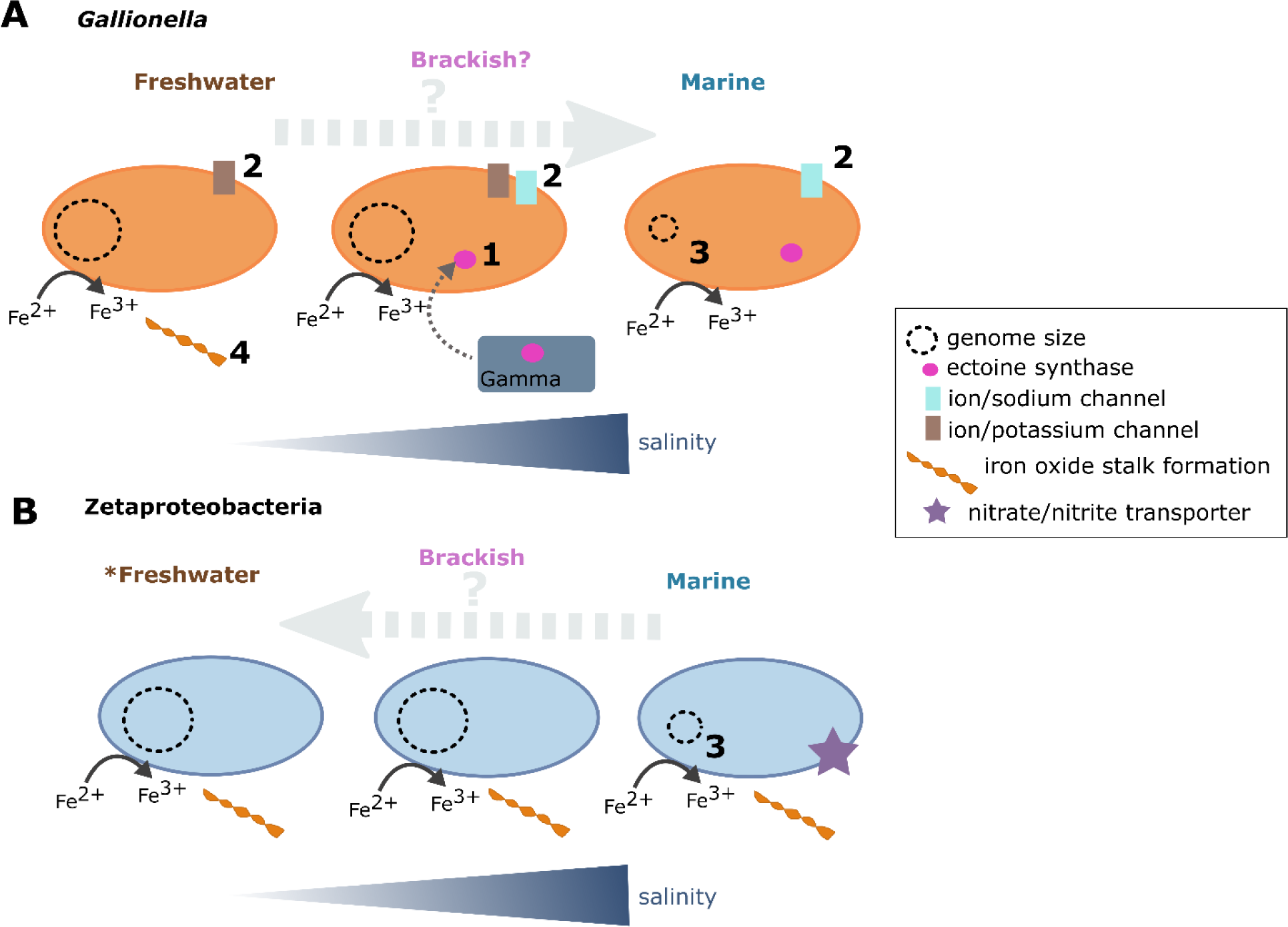
Model of adaptations in Gallionella (A) and Zetaproteobacteria (B) crossing the marine-freshwater barrier. The model is based on predicted genes in MAGs, involving adaptations discussed in this paper. 1) Ectoine synthase in marine Gallionella MAGs (possible horizontal gene transfer with Gammaproteobacteria). 2) Ion/potassium channels in freshwater microorganisms and ion/sodium channels in marine microorganisms. 3) Smaller genomes in marine microorganisms. 4) Formation of Fe(III) oxyhydroxide stalks based on putative stalk formation genes found in some representatives of Gallionella and Mariprofundus, however not in any known marine Gallionella MAGs. *Some MAGs are saline terrestrial, since in some terrestrial hot springs, salinity of 9-14 ppt was measured.

Marine *Gallionella* and Zetaproteobacteria genomes generally fall on the lower end of observed genome sizes compared to their freshwater and terrestrial counterparts, aligning with the globally observed trend across salinities (21, 22, 93). We stress that sampling of more genomes is required to statistically conclude that marine genomes of both *Gallionella* and Zetaproteobacteria are measurably smaller and/or went through genome streamlining. Distinctions in genome size are thought to arise from differences in the physicochemical environment. As freshwater habitats generally experience more rapid and pronounced fluctuations of physicochemical conditions compared to larger marine bodies of water, a larger genome of freshwater microorganisms enables them to be more flexible to adapt (19, 93).

Proteomes have previously shown more acidic values of isoelectric points (pI) in microbes exposed to marine environments and higher salinities than in freshwater and brackish microorganisms (23, 25, 93, 120, 121). Transitions between marine and freshwater environments are proposed to require long evolutionary time (23) and difficult to attribute solely to horizontal gene transfer (1). The isoelectric point values observed in *Gallionella* predicted proteomes showed no significant variations across the marine-freshwater divide, indicating that the proteome did not undergo extensive changes in its amino acid compositions. However, values and power analysis indicate that sampling of more genomes is required to draw concrete conclusions on whether larger proteome changes were involved in *Gallionella* salinity transitions. Nevertheless, this observation raises a strong argument that *Gallionella* transitions to the marine environment did not require wider genomic adaptations but rather involved horizontal gene transfer and occurred on a shorter evolutionary time scale.

### 3.4 Adaptations of Zetaproteobacteria to freshwater environments are less obvious

Presence of Zetaproteobacteria in certain terrestrial springs is possibly explained by elevated salinity levels (73), with examples of salinity between 9 and 14 ppt (78–80). The grouping may therefore not completely accurately reflect the actual physicochemical environment of the microorganisms, often excluding salinity. Due to the absence of salinity measurements for several terrestrial environments characterized and assumed as “freshwater”-associated (75, 83), it remains uncertain whether Zetaproteobacteria have the capability to inhabit “true” freshwater environments with low salinity. Patterns for genome and proteome adaptation in Zetaproteobacteria MAGs were however less intelligible than for *Gallionella.* Marine Zetaproteobacteria genomes group with those from seemingly freshwater environments with unknown salt concentrations, making a thorough analysis reliant on quality metadata on environmental parameters. Still, Zetaproteobacteria have shown environment differences in osmoregulation-related adaptations such as differences in genes for compatible solutes and involved in transport. In addition, nitrate assimilation genes present exclusively in marine Zetaproteobacteria genomes (Fig 7) suggest certain differences in nutrient uptake depending on the environment, argued to likely correspond to differences in availability of nitrate (41). While absent from other environments, nitrogen fixation genes were also previously observed in Zetaproteobacteria from hot springs (77). Absence of nitrate and nitrite reductases in freshwater Zetaproteobacteria suggests an aerobic lifestyle, with nitrate reduction potentially not playing a big role in freshwater Zetaproteobacteria. Since hydrothermal vent fluids contain large amounts of heavy metals (122, 123) which iron oxides adsorb (124–126), FeOB at hydrothermal vents could experience more heavy metal stress than in other environments. This would lead to more heavy metal resistance genes encoded in the genomes, such as mercury resistance as observed in hydrothermal Zetaproteobacteria (Fig S4).

### 3.5 Conclusion

Metagenome-resolved genomes of *Gallionella* were retrieved from deep sea hydrothermal vents, despite belonging to the previously recognized freshwater genus of iron-oxidizing bacteria. We provide further evidence that marine *Gallionella* can co-exist with Zetaproteobacteria FeOB and that they share an iron-oxidizing niche. Functional enrichment analyses showed several adaptations to an environment with elevated salinity. Even though only a few MAGs are available to date, our findings contribute to a better understanding of the diversification of FeOB, shedding light on their roles in the environment and their strategies for adapting to different conditions crossing the marine-freshwater barrier, including hydrothermal vents and freshwater habitats.

## 4 Materials and Methods

Details on methods used and reported in this study are found in Supplementary Material 2, deposited in a Zenodo repository.

### 4.0 Sample collection and processing

Samples were collected at two hydrothermal vent fields at the Arctic Mid-Ocean Ridges (AMOR). Samples were collected using an Ægir6000 remotely operating vehicle (ROV) on board the R/V G.O. SARS in July 2011, July 2012, and June 2019. The ROV was equipped with a biosyringe, a hydraulic sampling cylinder, linked to the ROV’s manipulator arm. Iron oxide deposit (FeDeposit) and outer chimney wall (ChimneyWall) were collected at Fåvne vent field (84, 127–129) at approx. 3036 m below sea level (Fig 2 C, Supplementary Material Table S1). Fe mat samples (Mat1, Mat2_I, Mat2_II) were collected from the Rift valley at Troll Wall vent field, one of Jan Mayen vent fields (55, 56, 130) at approx. 615 m below sea level (Fig 2 B, Supplementary Material 1 Table S1). Temperature readings were obtained using a temperature probe. The retrieved samples were centrifuged for 5 minutes at 6000 rcf, and the supernatant was discarded. Aliquots were frozen rapidly using liquid nitrogen and kept at –80°C until processing. Samples for scanning electron microscopy were fixed in a solution of 2.5% glutaraldehyde and kept at 4 °C until further processing. Scanning electron microscopy (SEM), elemental composition analysis and genome-resolved metagenomics using Illumina NovaSeq sequencing were performed as in a previous study (84).

## 5 Data availability

All MAGs in the study were deposited in NCBI and accession numbers with associated BioProject and BioSamples with corresponding metadata are listed in Supplementary Material 1 Table S1, Table S2, Table S3 and Table S4. Supplementary material is available online, deposited in a <u>Zenodo repository</u>.

## 7 Author Contributions

Author contributions are assigned using CRediT roles.

**Petra Hribovšek**: Conceptualization (lead), data curation (equal), formal analysis (lead), investigation (lead), methodology (lead), visualization (lead), writing – original draft (lead), writing – review & editing (lead). **Emily Olesin Denny**: Data curation (lead), writing – original draft (supporting). **Håkon Dahle**: Writing – original draft (supporting). **Achim Mall**: Data curation (equal). **Ida Helene Steen**: Conceptualization (lead), formal analysis (supporting), funding acquisition (lead), investigation (supporting), methodology (supporting), supervision (lead), writing – original draft (supporting), writing – review & editing (supporting). **Runar Stokke**: Conceptualization (lead), data curation (lead), formal analysis (supporting), funding acquisition (lead), investigation (supporting), methodology (lead), supervision (lead), project administration (lead), writing – original draft (supporting), writing – review & editing (supporting).

## 8 Acknowledgments

We thank the ROV Ægir6000 operator team and the R/V G.O. SARS crew for assistance during sampling, and Steffen L. Jørgensen for organizing the 2019 research cruise to the Fåvne vent field. We thank Irene Heggstad at ELMILAB for assistance with scanning electron microscopy. We are thankful to Eoghan P. Reeves for discussions on salinity. The computations associated to taxonomic classification and annotation of genes were performed on resources provided by Sigma2 – the National Infrastructure for High Performance Computing and Data Storage in Norway.

